# A bootstrap model comparison test for identifying genes with context-specific patterns of genetic regulation

**DOI:** 10.1101/2023.03.06.531446

**Authors:** Mykhaylo M. Malakhov, Ben Dai, Xiaotong T. Shen, Wei Pan

## Abstract

Understanding how genetic variation affects gene expression is essential for a complete picture of the functional pathways that give rise to complex traits. Although numerous studies have established that many genes are differentially expressed in distinct human tissues and cell types, no tools exist for identifying the genes whose expression is differentially regulated. Here we introduce DRAB (Differential Regulation Analysis by Bootstrapping), a gene-based method for testing whether patterns of genetic regulation are significantly different between tissues or other biological contexts. DRAB first leverages the elastic net to learn context-specific models of local genetic regulation and then applies a novel bootstrap-based model comparison test to check their equivalency. Unlike previous model comparison tests, our proposed approach can determine whether population-level models have equal predictive performance by accounting for the variability of feature selection and model training. We validated DRAB on mRNA expression data from a variety of human tissues in the Genotype-Tissue Expression (GTEx) Project. DRAB yielded biologically reasonable results and had sufficient power to detect genes with tissue-specific regulatory profiles while effectively controlling false positives. By providing a framework that facilitates the prioritization of differentially regulated genes, our study enables future discoveries on the genetic architecture of molecular phenotypes.

## 1. Introduction

Differential gene expression analysis is among the most common techniques in computational biology. It is used to compare the RNA transcript abundance of a gene between two different biological contexts or experimental conditions, and to determine whether the difference is greater than what would be expected due to natural random variation. A wide variety of statistical tools have been developed for differential expression analysis, from simple *t*-tests to sophisticated Bayesian approaches (Pan (2002); Bullard et al. (2010); Soneson and Delorenzi (2013); Seyednasrollah, Laiho and Elo (2015)). When applied to tissue-specific gene expression measurements, such as those available from the Genotype-Tissue Expression (GTEx) Project (Lonsdale et al. (2013); The GTEx Consortium (2020)), differential expression analysis can identify a set of genes whose expression is significantly different between two given tissues.

Gene expression studies can also provide insight into the genetic architecture of complex human diseases. Indeed, many recent approaches have sought to uncover causal loci by considering the functional pathways through which genes affect traits (Gallagher and Chen-Plotkin (2018); Wong et al. (2021)). A notable example is the transcriptome-wide association study (TWAS) framework, which prioritizes disease-relevant genes by testing for association with genetically regulated expression levels (Gamazon et al. (2015); Gusev et al. (2016)). TWAS has been applied in over a thousand studies and has led to a number of innovative methodological extensions (Barbeira et al. (2018); Hu et al. (2019); Yang et al. (2020); Lin et al. (2022)). The GTEx Project, for instance, utilized TWAS and related methods to characterize genetic effects on the transcriptome across human tissues and to link these regulatory mechanisms to trait and disease associations (The GTEx Consortium (2020); Barbeira et al. (2021)).

Yet despite the ubiquity of differential expression analysis and the importance of understanding how genetic regulation induces differences in gene expression, a fundamental problem remains poorly studied: when does the genetic regulation of expression differ between various tissue and cellular contexts?

This question is challenging to answer for at least two reasons. First, differential regulation is distinct from differential expression. For some genes, differences in expression levels are driven by fluctuations in tissue or cell microenvironment rather than by differences in networks of genetic regulatory activity. As a result, the presence of differential gene expression does not necessarily imply that context-specific patterns of genetic regulation exist. By the same reasoning, differentially regulated genes may not actually be differentially expressed. Second, the genetic architecture of gene expression is complex. Whereas context-specific gene expression levels can be easily quantified, genetic regulatory profiles might differ between biological contexts in a myriad ways. For example, consider two human tissues: different variants might be responsible for regulating gene expression in each tissue, or the same variants might have different effect sizes, or one tissue might be enriched in expression quantitative trait loci (eQTLs) relative to the other, etc. Therefore, patterns of genetic regulation cannot be summarized by a single index or compared using classical statistical tests.

In this paper we introduce DRAB (Differential Regulation Analysis by Bootstrapping), a statistical method for determining whether gene-level genetic regulation is significantly different between two tissues or other biological contexts. The premise of our approach is to train penalized regression models that capture the context-specific effects of local genetic variation on gene expression, and then to statistically test whether those models have equal predictive performance on an independent test set. During the training process each model learns the particular regulatory patterns present in the context that it is trained on, and so testing for model equivalency effectively tests whether the genetic regulation of expression is equivalent between those contexts. To perform such a test without conditioning on a specific pair of trained models, we develop a novel bootstrap-based model comparison framework. The DRAB test accounts for the uncertainty of performing feature selection and model fitting on randomly sampled training sets, allowing us to interpret it as a test of equivalency between population-level models of genetic regulation. As such, DRAB directly addresses both of the challenges outlined above. To our knowledge this is the first statistical method for identifying genes with tissue-specific (or other context-specific) patterns of genetic regulation.

We demonstrate DRAB by applying it to autosomal genotypes and mRNA gene expression data from 23 selected human tissues in the GTEx v8 data release (The GTEx Consortium (2020)). We show that our method is able to control type I error and has sufficient power to detect tissue-specific differences in genetic regulatory profiles, while conventional model comparison tests yield many spurious results. By evaluating a broad array of tissue pairs, we confirm that the genetic regulation of gene expression is more similar in biologically related tissues (e.g. different brain tissues) than in unrelated tissue pairs (e.g. blood and brain tissues). In particular, we identify sets of genes whose expression is differentially regulated between blood and various brain regions, between distinct nervous system tissues, and between a variety of other selected tissue pairs. To facilitate further discovery we make an R implementation of DRAB available on GitHub, along with supporting bash scripts for data processing (https://github.com/MykMal/drab).

## 2. Methods

### 2.1. Overview of DRAB

Consider a gene *g*, and let *A* and *B* denote two tissues, disease states, or other biological sample types. The purpose of DRAB is to test the null hypothesis that the *cis*-genetic regulation of *g* is equivalent in context *A* and context *B*. To do this, DRAB first trains elastic net models in each context to predict the expression of *g* from single nucleotide polymorphisms (SNPs) located near the gene. The training algorithm simultaneously performs feature selection and model fitting, so that the resulting models capture context-specific patterns of genetic regulation in *cis* for gene *g*. Then DRAB tests whether those context-specific models are statistically equivalent using a novel model comparison test on an independent test set. Our proposed model comparison test differs from previous approaches in that it accounts for the uncertainty of learning a regression function on randomly sampled training sets, which we estimate through bootstrap resampling. The two stages of DRAB are visualized in Figure 1 and detailed below. In practice, DRAB automatically repeats this process for each gene on a user-specified list of genes. Note that individual-level genotype and gene expression data are required.

**FIG 1.**
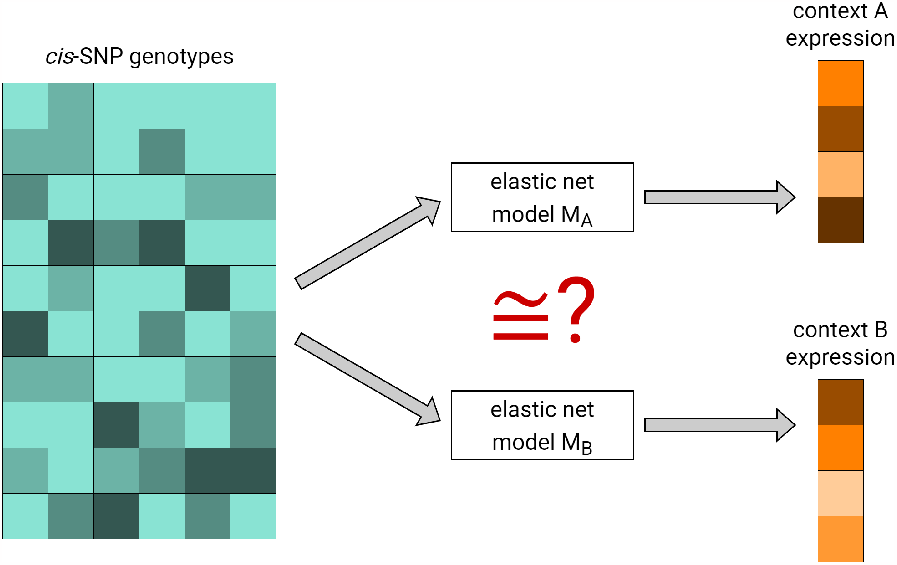
The DRAB method is comprised of two stages. First, elastic net models are trained to predict gene expression from cis-SNPs in each context. Second, the context-specific models are tested for statistical equivalency at the population level using a bootstrap-based model comparison test on an independent test set.

### 2.2. Stage 0: data preprocessing

Let *D*_*A*_ be the training set for context *A, D*_*B*_ be the training set for context *B*, and *D*_*T*_ be the test set. In subsequent stages of DRAB, *D*_*A*_ and *D*_*B*_ are used for training context-specific models of genetic regulation, while *D*_*T*_ is used for comparing the two models’ predictive performance. To ensure that comparisons of predictive performance are unbiased, we require that *D*_*T*_ ∩ (*D*_*A*_ ∪*D*_*B*_) =. In addition, we assume that |*D*_*A*_| = |*D*_*B*_| to avoid the possibility of one fitted model being less stable than the other, which would decrease the power of any model comparison test. That is, the two training sets must have the same cardinality and each be independent from the test set. By default, our implementation of DRAB will automatically create the largest possible training sets *D*_*A*_, *D*_*B*_ from the provided input data and then set aside the remaining data for the test set *D*_*T*_. These defaults are intended to maximize efficiency, but our code also provides the option of specifying custom training and test data splits.

Finally, in this paper we assume that both data sets are drawn from the same population. Although DRAB can also be used for studies where context *A* was measured in a distinct population than context *B*, this would change the interpretation of results. It is well-known that the prediction accuracy of transcriptome imputation models greatly suffers when the training and test populations differ in terms of allele frequency or linkage disequilibrium (LD) patterns (Keys et al. (2020); Okoro et al. (2021)). Since DRAB relies on prediction error to determine model equivalency, a discrepancy in population ancestry between contexts may lead to some genes being identified as significant due to ancestry-specific differences in genetic regulation, rather than due to context-specific differences. To avoid such confounding and ensure that our results solely reflect tissue-specific genetic regulatory patterns, we restricted our real data application to individuals with European genetic ancestry.

DRAB begins by extracting SNPs that are located within the *cis*-locus of *g*, which we define as ranging from 500 kilobase pairs upstream of the gene’s transcription start site to 500 kilobase pairs downstream of its transcription end site. We refer to the variants within this region as *cis*-SNPs. Next, the expression levels of *g* and the genotypes of each *cis*-SNP are adjusted for a set of user-specified covariates. Covariates are regressed out separately on each of the sets *D*_*A*_, *D*_*B*_, and *D*_*T*_ to preserve their independence. Then we center and standardize the expression residuals, as well as the residuals of each *cis*-SNP, to have a mean of zero and a variance of one. As a final quality-control step, we remove any monomorphic *cis*-SNPs. The normalized residuals are subsequently used as the expression and genotype values in the main stages of DRAB.

### 2.3. Stage 1: model training

DRAB independently trains two parametric machine learning models: model *M*_*A*_ and model *M*_*B*_. Model *M*_*A*_ is trained to predict the (adjusted) expression levels of gene *g* in context *A*, using data from the individuals in training set *D*_*A*_. Similarly, model *M*_*B*_ is trained to predict the (adjusted) expression levels of gene *g* in context *B*, using data from the individuals in training set *D*_*B*_. Both models are given the same set of (adjusted) *cis*-SNPs as features, but differ in their response variables and training sets.

In particular, we use Gaussian family elastic net models (Zou and Hastie (2005)). The elastic net is a penalized regression method that linearly combines the *L*_1_ norm from lasso regression (Tibshirani (1996)) and the *L*_2_ norm from ridge regression (Hoerl and Kennard (1970)). Consequently, the elastic net simultaneously performs both regularization and variable selection, leading to its popularity in applications that require learning sparse models from high-dimensional data with correlated features. Given genotype dosage vectors *x*_*i*_ ∈ R^*p*^ and gene expression measurements *y*_*i*_ ∈ R for individuals *i* = 1, …, *N*, the elastic net minimizes the objective function

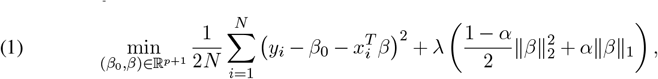

where *λ* ≥ 0 is a complexity parameter and 0 ≤ *α* ≤ 1 controls the relative weight of each penalty. We fix the mixing parameter *α* at *α* = 0.5, which tends to either select or leave out entire groups of correlated features (Zou and Hastie (2005)). As a result, *M*_*A*_ and *M*_*B*_ each select the groups of *cis*-eQTLs that are specific to the contexts they are trained on. By performing both feature selection and model fitting, the elastic net efficiently learns tissuespecific patterns of *cis*-genetic regulation.

To train the elastic net models, DRAB relies on the R package `glmnet` (Friedman, Hastie and Tibshirani (2010); Tibshirani et al. (2012)). The `glmnet` package dynamically computes a grid of values for the tuning parameter *λ*, ensuring that the grid covers the entire range of possible solutions. Then we use 5-fold cross validation to select the value of *λ* that yields the most parsimonious model whose cross-validated error is within one standard deviation of the minimum. DRAB utilizes this approach instead of selecting the best possible *λ* because sparse models tend to outperform more polygenic models at transcriptome prediction (Wheeler et al. (2016); Fryett, Morris and Cordell (2020)).

Note that other algorithms for regression and feature selection could be justifiably used here instead of a linear model with the elastic net penalty. Our proposed model comparison test relies on empirical distributions and can thus be performed with any parametric regression models. However, the elastic net algorithm offers several desirable properties that make it the most natural choice for our application. Most importantly, the elastic net selects sparse groups of context-specific SNPs and learns context-specific weights for each selected SNP. This makes DRAB robust to the high dimensionality of genetic data and the inherent complexity of LD between eQTLs involved in transcriptional regulation. Moreover, the elastic net is commonly used for transcriptome imputation in TWAS (Gamazon et al. (2015); Barbeira et al. (2018); Gillies et al. (2018); Wu et al. (2018); Guo et al. (2021)) and previous research has shown that it outperforms several other machine learning algorithms at this task (Okoro et al. (2021)).

### 2.4. Stage 2: testing model equivalency

#### 2.4.1. Definitions

Our goal is to conduct a model comparison test in order to determine whether the stage 1 models *M*_*A*_ and *M*_*B*_ are statistically equivalent at the population level with respect to some loss function *L*. Here we must carefully define the concept of “models” and what we mean when speaking of their equivalence. To distinguish the fitted stage 1 models from the theoretical population-level models, we will use the notation 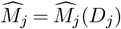 to signify the model trained on set *D*_*j*_ for *j* = *A, B*. In other words, *M*_*j*_ is the true function that captures the effect of genetic variation on gene expression for context *j*, while *M*_*j*_ is the model returned by some algorithm that attempts to learn this functional relationship based on training data *D*_*j*_. We assume that the trained models are consistent for the true models: 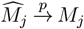 as |*D*_*j*_| → ∞ for *j* = *A, B*. That is, we assume that the finite-sample fitted models will converge in probability to the true population-level models as the training sample sizes increase to infinity.

We say that population-level models *M*_*A*_ and *M*_*B*_ are equivalent with respect to some loss function *L* when their prediction errors with respect to *L* are equal for all samples in the population. As such, the definition of equivalence depends on the choice of *L*: two models may be equivalent with respect to one loss function, but not equivalent with respect to another. Here we will exclusively consider the squared error loss, although our framework can be applied to any valid loss function. This formulation of model equivalence is conducive for developing a formal model comparison framework, because it enables us to determine whether models are equivalent by testing for equality between conditional means. Further details are provided in the subsequent sections.

Our definition of equivalence partitions the space of all models into a number of equivalence classes. Importantly, models that always have the same prediction accuracy with respect to *L* belong to the same equivalence class and cannot be distinguished even if they have distinct functional forms. Such situations may arise in practice due to LD between the genetic variants used as features. For example, suppose there is a single causal variant that affects gene expression, but it lies on a haplotype with other variants, and this block of variants is preserved throughout the population. Then the sparse causal model is not identifiable in terms of prediction loss from another model making use of other variants on the haplotype, such as one which distributes the effect uniformly across all variants on that haplotype. The definition of model equivalence and its implications must be considered when interpreting the results of DRAB or any other model comparison test.

Since the true models *M*_*A*_ and *M*_*B*_ are unknown, we must instead work with the fitted models 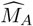 and 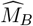. Let 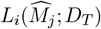 denote the squared prediction residual for model 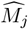 and individual *i* ∈ *D*_*T*_. Then the average difference in prediction errors between model 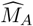 and model 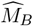 on the test set *D*_*T*_ is

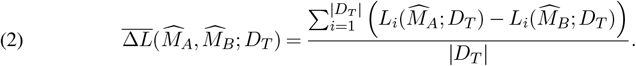

#### 2.4.2. Limitations of conditional approaches

The standard practice for model comparison is to evaluate both models on an independent test set and compare their prediction errors. When a formal decision is desired, one can test the null hypothesis that the prediction errors are equal. For regression learning tasks, in particular, a paired samples *t*-test on the squared residual errors is used. Here we review this approach and explain why it is inadequate for assessing whether context-specific models of genetic regulation are equivalent.

The paired samples *t*-test relies on three assumptions: prediction errors are “paired” between the two models, each pair is independent of the rest, and the pairwise differences are approximately normally distributed. The first assumption is satisfied because prediction errors for both models are computed on the same test set, so 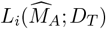 and 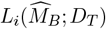 form a pair for each individual *i* ∈ *D*_*T*_. We follow the precedent in large-scale genetics studies of assuming that genotyped individuals are unrelated, which ensures that the second assumption is satisfied. Finally, the third assumption follows from the central limit theorem for a sufficiently large test set.

The paired samples *t*-test statistic is given by

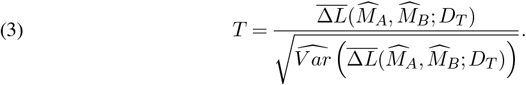

Variance can be estimated in the standard way as

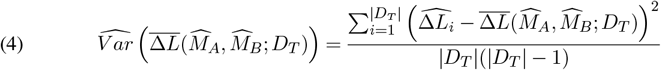

Where 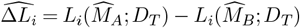. Under the null hypothesis that models 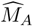 and 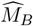 have equal predictive performance with respect to *L*, the statistic *T* follows a Student’s *t*-distribution with |*D*_*T*_| − 1 degrees of freedom.

Although this model comparison framework is commonly used in the statistical learning literature, it has a major limitation. By definition, the *t*-test described above determines whether the mean difference between prediction errors for model 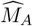 and model 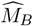 is zero. Importantly, it pertains to the fitted models *M*_*A*_, *M*_*B*_ rather than the theoretical populationlevel models 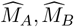. For some applications, conditioning on the trained models is satisfactory or even desired; but in our case ignoring the uncertainty associated with model training would lead to a greatly inflated false positive rate.

It is well known that models pick up finite-sample errors during the training process. For example, fitting the same linear regression model on two separate data sets will yield different predictor coefficients—even if both data sets are randomly sampled from a common population. The stochastic nature of data sampling becomes especially problematic when performing feature selection, since the same feature selection algorithm might choose different sets of features when applied to different random samples from the same population (Kalousis, Prados and Hilario (2007); Nogueira, Sechidis and Brown (2018)). The issue of model training uncertainty is further exacerbated in algorithms that have a stochastic component, such as cross validation. Such algorithms are known to select different sets of features when repeatedly trained on exactly the same data set (Cadiou and Slama (2021)). Therefore, the elastic net algorithm that we use in stage 1 suffers from instability both due to the uncertainty of tuning the *λ* hyper-parameter using cross validation and the uncertainty of estimating coefficients on randomly sampled finite training sets.

As a result of these dual uncertainties associated with performing feature selection and model fitting, the trained stage 1 models 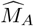 and 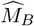 can be significantly different even when the theoretical models *M*_*A*_ and *M*_*B*_ are equivalent. The aim of DRAB is to compare the population-level patterns of genetic regulation between contexts *A* and *B*, so testing for equivalency between the trained models 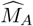 and 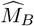 would lead to erroneous conclusions. In fact, we show in Section 3.2 that attempting to use the standard *t*-test for comparing models of genetic regulation yields unacceptable results.

#### 2.4.3. The DRAB model comparison test

To mitigate the problem of model training variability, we developed a new model comparison test based on the non-parametric bootstrap. Unlike the standard model comparison approach described above, our test explicitly accounts for the uncertainty of feature selection and model fitting. Hence, it can be interpreted as testing whether the theoretical population-level models, *M*_*A*_ and *M*_*B*_, are statistically equivalent. Our proposed test is inspired by recent work on hypothesis testing for black-box models (Wasserman, Ramdas and Balakrishnan (2020); Dai, Shen and Pan (2022)), but it is methodologically novel and conceptually distinct.

The key insight behind the DRAB model comparison test is that we avoid conditioning on the fitted models 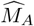 and 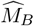, instead estimating what the paired samples *t*-test statistic would be under the true models of genetic regulation. Formally, we wish to test the null hypothesis

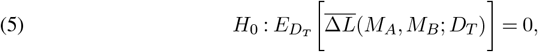

which states that the population means of the prediction errors of models *M*_*A*_, *M*_*B*_ are equal with respect to the loss function *L*. It is intuitive to imagine the following statistic for this test:

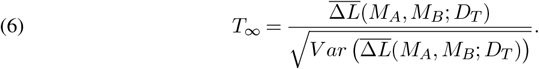

But the theoretical population-level models *M*_*A*_ and *M*_*B*_ are unknown by definition! We therefore resort to approximating the numerator of *T*_*∞*_ and its variance.

Since 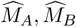 are consistent for *M*_*A*_, *M*_*B*_ by our earlier assumption, the naive estimator for 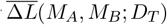 will yield good results as long as the training data sets are sufficiently large. Thus, we approximate the numerator of *T*_*∞*_ by substituting in the full-sample trained models:

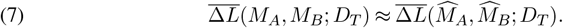

Now let 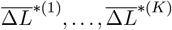 be *K* non-parametric bootstrapped replicates of 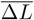 That is, for each *k* = 1, …, *K* we create the sets 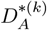 and 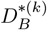 by randomly sampling with replacement from *D*_*A*_ and *D*_*B*_, respectively, such that 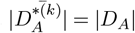 Then on each set 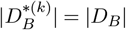 we train a model to predict expression in context *A*, and denote it by 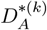. Similarly, on each set 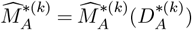 we train a model to predict expression in context *B*, and denote it by 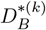. Finally, for each *k* we compute

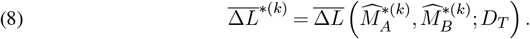

The resulting empirical distribution of 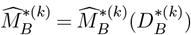 approximates the sampling distribution of 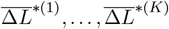 across all possible context-specific training sets (Efron and Tibshirani (1993)).

To estimate the variance of 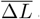(*M_A_;M_B_D_T_;*), we decompose it using the law of total variance. Observe that

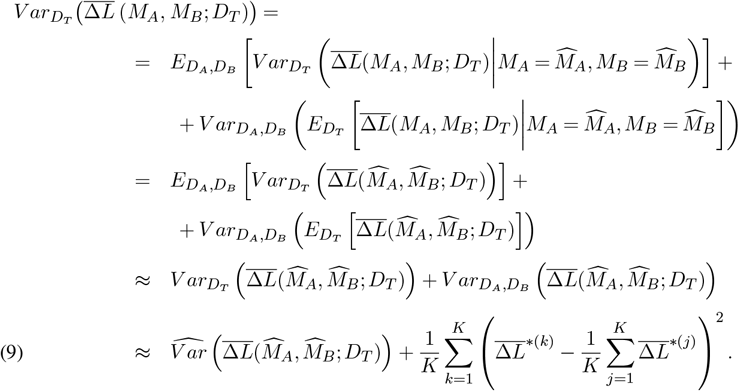

Note that in the last line we used the bootstrap distribution of 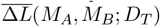 to approximate the variance of 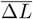 across all possible training sets.

This variance estimate can be interpreted as the sum of two sources of uncertainty. The first term accounts for the uncertainty of the estimator 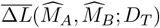 computed on the test set *D*_*T*_, so it will converge to zero as |*D*_*T*_ | *→ ∞*. The second term, on the other hand, accounts for the uncertainty of the estimated (i.e. trained) models 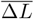 and 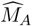, so it will converge to zero |*D*_*A*_ | *→ ∞*. and |*D*_*B*_ | *→ ∞*.only when *D*_*A*_ and *D*_*B*_. In other words, the first term pertains to the sampling variability of the test set *D*_*T*_ while the second term pertains to the sampling variabilities of the training sets *D*_*A*_ and *D*_*B*_. Including both sources of variance is the primary innovation of the DRAB test, which allows it to overcome the limitations of model comparison tests that condition on a specific pair of fitted models.

Putting everything together, our proposed DRAB test statistic is

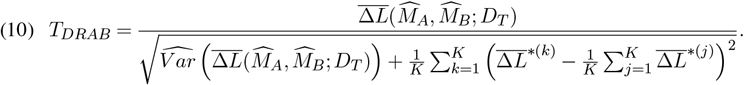

We claim that under the null hypothesis stated in Equation 5, the *T*_*DRAB*_ statistic approximately follows a Student’s *t*-distribution with |*D*_*T*_| − 1 degrees of freedom. Our proposed statistical test expressly accounts for the variability in estimating *M*_*A*_ and *M*_*B*_ through model training, and so it can be used to determine whether the true theoretical models are statistically equivalent. If the bootstrap-based DRAB test rejects the null hypothesis of model equivalency, we conclude that the patterns of genetic regulation for gene *g* are significantly different in contexts *A* and *B*.

## 3. Application

### 3.1. Data preparation

To validate DRAB and demonstrate its use, we applied it on data from the GTEx v8 release (The GTEx Consortium (2020)). In particular, we downloaded individual-level genetic variants as a Variant Call Format (VCF) file from the database of Genotypes and Phenotypes (dbGaP), under accession number phs000424.v8.p2. The VCF file was derived from whole-genome sequencing data as described in The GTEx Consortium (2020). Extensive sample quality control procedures, variant quality control procedures, and read-aware phasing had already been performed on the genotype data, leaving 838 individuals and 46,526,292 genomic sites. We also downloaded tissue-specific matrices of bulk gene expression from the GTEx Portal (https://gtexportal.org/home/datasets). The gene expression data had already been quantified, processed, and normalized as described in The GTEx Consortium (2020) and was reported in terms of Transcripts Per Million (TPM) units. Data for 49 human tissues was available, with the number of quantified genes ranging from 20,249 in whole blood to 35,054 in testis.

Prior to applying our method, we performed some additional processing on the genotype data. We first subset the individuals to only keep those of European descent. Next we removed all variants that are non-autosomal, as well as those that have at least one multi-character allele code or a nonstandard single-character allele code. Then we removed variants with missing genotype data and tested the remaining ones for Hardy-Weinberg equilibrium (HWE). In particular, we used the mid-p method to test for HWE (Wigginton, Cutler and Abecasis (2005); Graffelman and Moreno (2013)) and discarded SNPs with *P* < 10^*−*6^. Lastly, we filtered out SNPs with minor allele frequency (MAF) below 0.01. After all of these steps, 715 individuals and 8,042,975 genomic sites remained. All genetic quality control steps were conducted in PLINK 1.90 (Chang et al. (2015); Purcell and Chang) and the resulting files were imported into R through the `BEDMatrix` package (Grueneberg and de los Campos (2019)).

DRAB supports covariate adjustment with any user-specified covariates. In this study we included the pre-computed European covariates that are available from the GTEx Portal. Namely, we downloaded and included the top 5 genotyping principal components, a set of latent covariates identified using the Probabilistic Estimation of Expression Residuals (PEER) method, and indicators for sequencing platform (Illumina HiSeq 2000 or HiSeq X), sequencing protocol (PCR-based or PCR-free), and sex. For details on the definitions of these covariates, see The GTEx Consortium (2020).

### 3.2. The DRAB test accounts for model training uncertainty

To demonstrate the advantages of our proposed bootstrap model comparison test over conditional tests, we evaluated the performance of both approaches on a single-tissue negative control. The negative control scenario was created by partitioning the set of individuals for whom gene expression measurements are available in whole blood tissue (*n* = 558) into three equal-sized subsets. Two of these subsets were treated as the training sets *D*_*A*_ and *D*_*B*_, while the remaining one was treated as the test set *D*_*T*_. Note that since the three subsets are nonoverlapping and have equal sizes, they satisfy the independence and cardinality assumptions stated in Section 2.2. We then applied DRAB to these data sets. Additionally, we created a conditional version of the DRAB framework by replacing our bootstrap model comparison test with a paired samples *t*-test on the model prediction errors, as detailed in Section 2.4.2. The “conditional DRAB” test was also applied to the same data. We performed both of these differential regulation tests on each autosomal protein-coding gene, as defined by the GENCODE 26 transcript model provided on the GTEx Portal (The GTEx Consortium (2020)).

By construction, our negative control represents the scenario where context *A* and context *B* are sampled from the same population. Consequently, the patterns of genetic regulation are equivalent at the population level between the two contexts. If the context-specific models select different sets of eQTLs with differing effect sizes, then we know that those differences are only due to natural random variation and the instability of the learning algorithm. Ideally, a differential regulation analysis test should not identify any significant genes between the two subsets.

Figure 2 displays the per-chromosome results from our proposed DRAB test as well as from the analogous conditional test. The quantile-quantile (QQ) plots compare the observed distribution of *P* -values with their expected distribution under the null hypothesis of no differences in genetic regulatory patterns between the two subsets. Since we intentionally constructed the training data to ensure that the null hypothesis is true, the observed − log_10_(*P*) values on each QQ plot should mostly lie within the 95% confidence band surrounding the identity line (shown in gray). However, this is clearly not the case for the conditional model comparison test. When the paired samples *t*-test is used to determine model equivalency, we observe extreme inflation of the − log_10_(*P*) values. In fact, most of the *P* -values returned by the conditional test lie outside of the 95% confidence bands. Even after a Bonferroni correction for multiple testing, the conditional model comparison framework reported 19 transcriptome-wide significant genes. Clearly, this is an unacceptably high type I error rate for ascertaining the presence of population-level differences in genetic regulation.

**FIG 2.**
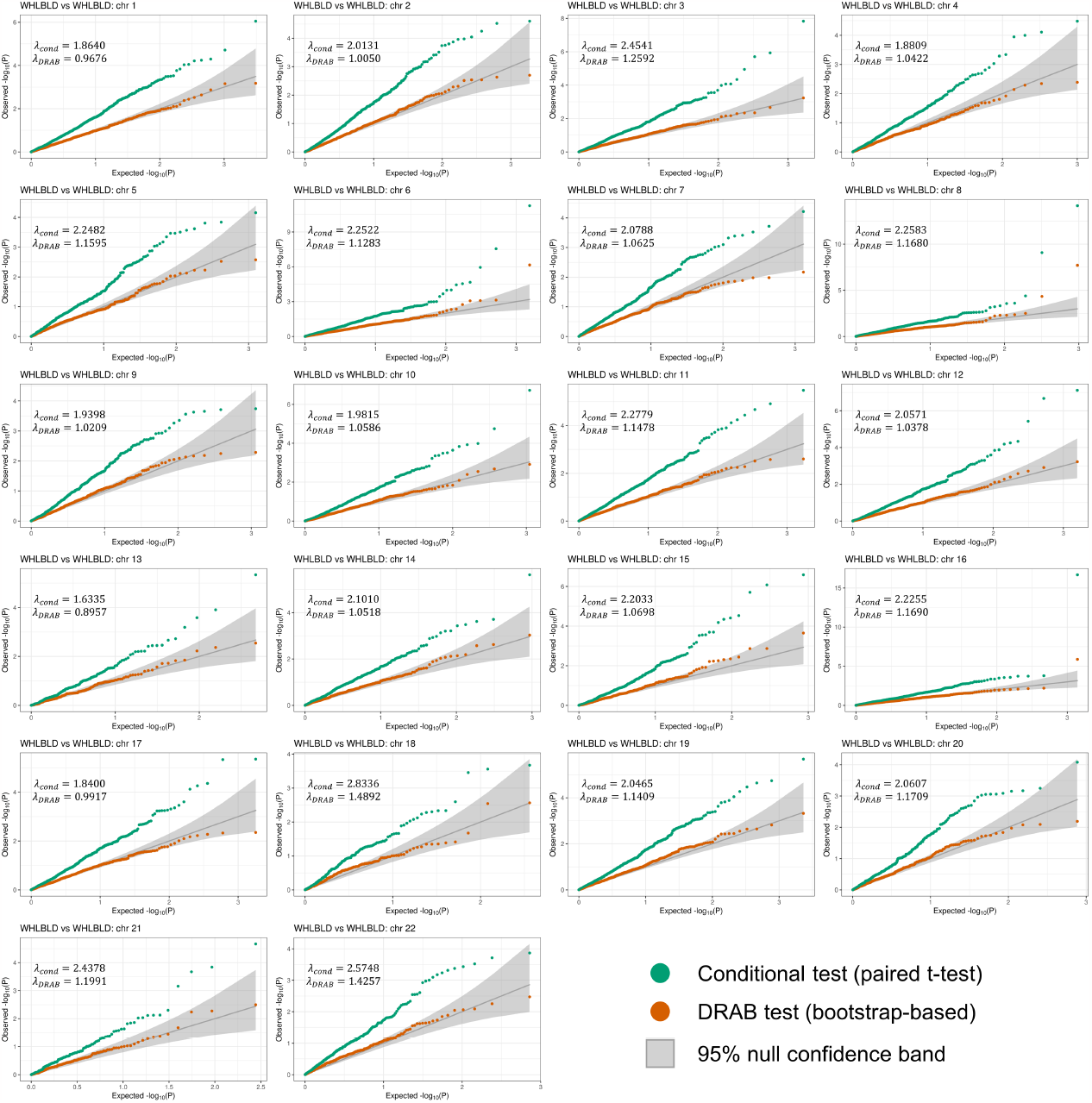
Per-chromosome quantile-quantile (QQ) plots of P -values from differential regulation tests on whole blood (WHLBLD) data. The full data set for whole blood was partitioned into three equal-sized subsets, with two subsets used for training transcriptome prediction models and the third subset used for testing. All autosomal proteincoding genes were included in the analysis. Orange points denote − log_10_(P) values from the bootstrap-based DRAB test, while green points denote − log_10_(P) values from a paired t-test on the model prediction errors. The theoretical null distribution (assuming no subset-specific genetic regulation) is plotted as a solid gray line, flanked by 95% confidence bands. λ_cond_ and λ_DRAB_ are genomic control lambdas estimated for the conditional t-test results and the DRAB test results, respectively.

Our proposed bootstrap model comparison test, on the other hand, was able to effectively control the false positive rate. Notice that all of the DRAB *P* -values lie within the 95% confidence bands of Figure 2 (with only a couple minor exceptions, likely due to correlation between genes). Although the DRAB test is sometimes slightly conservative (e.g. on chromosome 17) or slightly liberal (e.g. on chromosome 15), overall it eliminated the hugely inflated type I error rates that were observed with a conditional test. This is also evident by considering the genomic control lambda for each test, which we computed using the robust median estimator. Across all chromosomes, the conditional *t*-test exhibited an extremely high inflation of *λ*_*cond*_ = 2.0931, whereas the DRAB test results were only slightly inflated with *λ*_*DRAB*_ = 1.0967. These results demonstrate that our proposed model comparison test is able to account for the variability of feature selection and model training, enabling rigorous comparisons of regression models at the population level.

Surprisingly, DRAB was able to effectively control false positives with relatively few bootstrap iterations. For all of the figures in this paper we used *K* = 50 bootstrap iterations, a much lower number than the hundreds or thousands of iterations usually recommended for the non-parametric bootstrap. This reduces the number of times that models need to be re-fit, making our framework computationally feasible to implement in real-world settings. When running on one core of a third-generation AMD EPYC 7763 processor with *K* = 50 bootstrap iterations, DRAB took on average 1 minute 2 seconds per gene and utilized 543 MB of RAM given data with training sample sizes of 135 and a test set sample size of 423. Larger training set sample sizes, and to a lesser extent larger test set sample sizes, will increase runtime and memory usage. For example, given data with training sample sizes of 438 and a test set sample size of 120, DRAB took on average 1 minute 46 seconds per gene and utilized 755 MB of RAM. Moreover, we found that DRAB can yield acceptable results in the negative control scenario with as few as *K* = 10 bootstrap iterations, but results become unstable when the number of iterations is low.

### 3.3. DRAB is robust to diverse simulation settings

We conducted simulation studies to assess how sample overlap between the training sets, choice of tissue for the test set, and sample size affect the performance of DRAB. First, we trained transcriptome imputation models for whole blood and brain cortex as described in Section 2.3. From the genes whose models had nonzero weights in both tissues, we randomly selected 50 genes that DRAB identified as being differentially regulated (*P <* 0.05, Bonferroni corrected) in our real data application (see Section 3.4). The weights from these models were then used to predict gene expression levels in whole blood and brain cortex for every genotyped individual of European ancestry in the GTEx v8 data set (*n* = 715). Finally, we added independent Gaussian noise to the predicted expression levels to represent non-genetic effects on gene expression. The mean of the noise was fixed at zero and the variance was set to the residual variance from the model training step.

Our results demonstrate that sample overlap and choice of tissue for the test set have no impact on the type I error of DRAB. We varied the amount of sample overlap between *D*_*A*_ and *D*_*B*_ from 0% to 100% while keeping the training sample sizes fixed at *n* = 238 for each training set. Moreover, for each of the overlap scenarios we considered testing on either simulated brain cortex expression data or simulated whole blood expression data, keeping the test set sample size fixed at *n* = 239 and ensuring that it does not overlap with the training sets. We found that DRAB did not identify any false positives in null comparisons across all simulations, and the QQ plots remained qualitatively identical. The QQ plots for all 20 simulation setups are available in the Supplementary Material (Malakhov et al. (2023)).

Similarly, our simulation results show that sample overlap between the two training sets has no impact on the power of DRAB. We again varied the amount of sample overlap between *D*_*A*_ and *D*_*B*_ from 0% to 100% while keeping training and test sample sizes fixed. Across all simulation scenarios, DRAB *P* -values remained largely the same and no systematic change was observed as sample overlap increased. This was true regardless of whether whole blood or brain cortex was used as the test set tissue. See the Supplementary Material (Malakhov et al. (2023)) for scatter plots from all simulation setups. However, we observed that the choice of tissue for the test set can affect the power of DRAB. Tissues with greater variation in gene expression levels across individuals enable DRAB to detect more subtle differences in genetic regulatory patterns, since the test relies on a comparison between predicted and measured expression on the test set. In the extreme case, a tissue with constant expression for the given gene is useless as a test tissue and will result in zero power. In our simulations, whole blood expression was significantly better imputed than brain cortex expression, and as a result the simulated expression levels for blood captured more variation. This led to somewhat smaller *P* -values when simulated whole blood expression was used for the test set, as shown in the Supplementary Material (Malakhov et al. (2023)).

Finally, we assessed how training and test set sample sizes affect the power of DRAB to detect differentially regulated genes. Scatter plots comparing DRAB *P* -values from tests performed with different training and test set sample sizes are provided in the Supplementary Material (Malakhov et al. (2023)). We varied the test set sample size from *n* = 239 individuals to *n* = 30 individuals while keeping the training sample sizes fixed at *n* = 238 individuals each. As expected, the power of DRAB decreased with decreasing test set sample sizes. While results with |*D*_*T*_| = 179 and |*D*_*T*_| = 239 were comparable, *P* -values from tests with |*D*_*T*_| = 120 and below were systematically larger than *P* -values from tests with |*D*_*T*_| = 239. Power likewise decreased as we lowered the training set sample sizes from *n* = 328 to *n* = 30 while keeping the test set sample size fixed at |*D*_*T*_| = 239, especially when the sample size dropped below 100. Moreover, we observed that the test statistic became very unstable at the lowest training set sizes, indicating that insufficient data was available to obtain good estimates of the model weights. In general, sample size is a crucial factor for determining the power of any statistical test, and DRAB is not immune from such considerations.

### 3.4. Biologically related tissues share more regulatory patterns

Having verified that DRAB is able to control the type I error rate and is robust to sample overlap, we next assessed its ability to detect tissue-specific patterns of genetic regulation between distinct tissues. We applied DRAB to 19 pairs comprised of 23 selected human tissues from the GTEx v8 data release. Since running DRAB on all possible pairings of the 49 tissues available in GTEx would have been computationally prohibitive, we aimed to select tissue pairs that represent a broad spectrum of tissues and tissue relationships. In particular, we made sure to include comparisons between whole blood and various nervous system tissues because whole blood is the most commonly used bulk tissue in TWAS, owing to its relative ease of procurement. However, we suspected that blood might not be representative of genetic regulatory profiles in highly specialized brain tissues. We also tested for differential regulation between multiple brain regions. Finally, we chose six miscellaneous tissue pairs to further illustrate how extensively the genetic architecture of gene expression can vary between tissue types. The full list of considered tissues, along with their abbreviations and sample sizes, is displayed in Table 1.

**TABLE 1.**
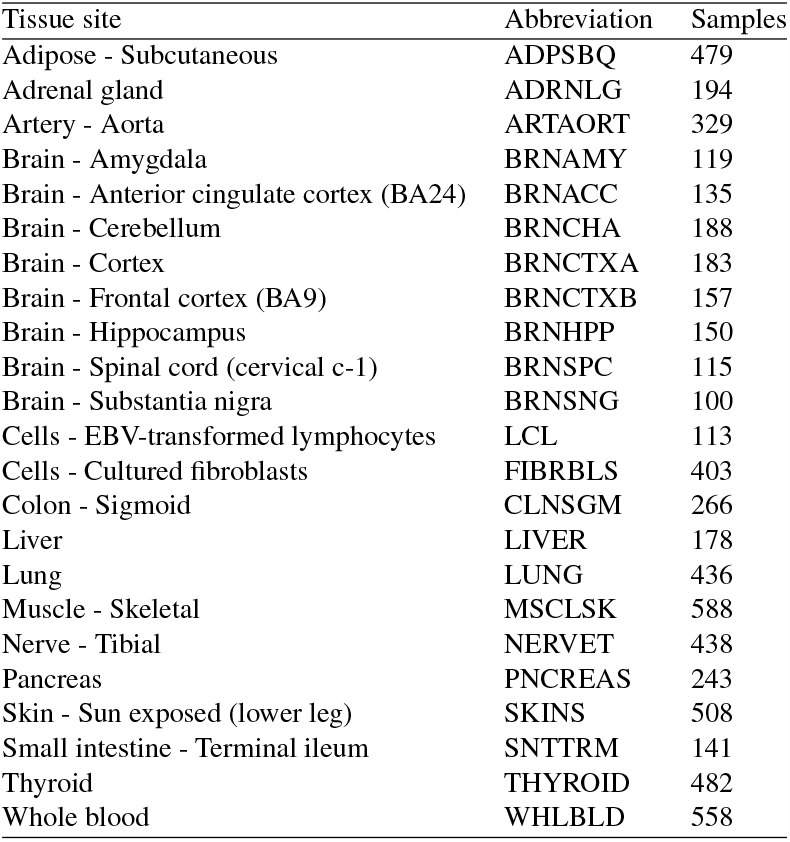
Full list of considered tissues.

Our results demonstrate that DRAB does have sufficient power to detect genes with tissuespecific patterns of genetic regulation in a real-world setting. Figure 3 shows Manhattan plots of DRAB test results for all autosomal protein-coding genes in three tissue pairs. The tissues in each of these pairs have markedly different biological function and structure; for example, blood is entirely unlike the hippocampus. Therefore, DRAB should be able to identify a large number of differentially regulated genes in each of these tissue pairs. The DRAB test was indeed able to do so. After a Bonferroni correction for multiple testing, 99 genes were transcriptome-wide significant between brain cortex and tibial nerve, 84 genes were transcriptome-wide significant between lung and aorta, and 172 genes were transcriptome-wide significant between whole blood and brain hippocampus. Figure 3 also displays a bar chart with the counts of significant protein-coding genes for each of the tissue pairs that we considered, while QQ plots with *P* -values from every pair are available in the Supplementary Material (Malakhov et al. (2023)).

**FIG 3.**
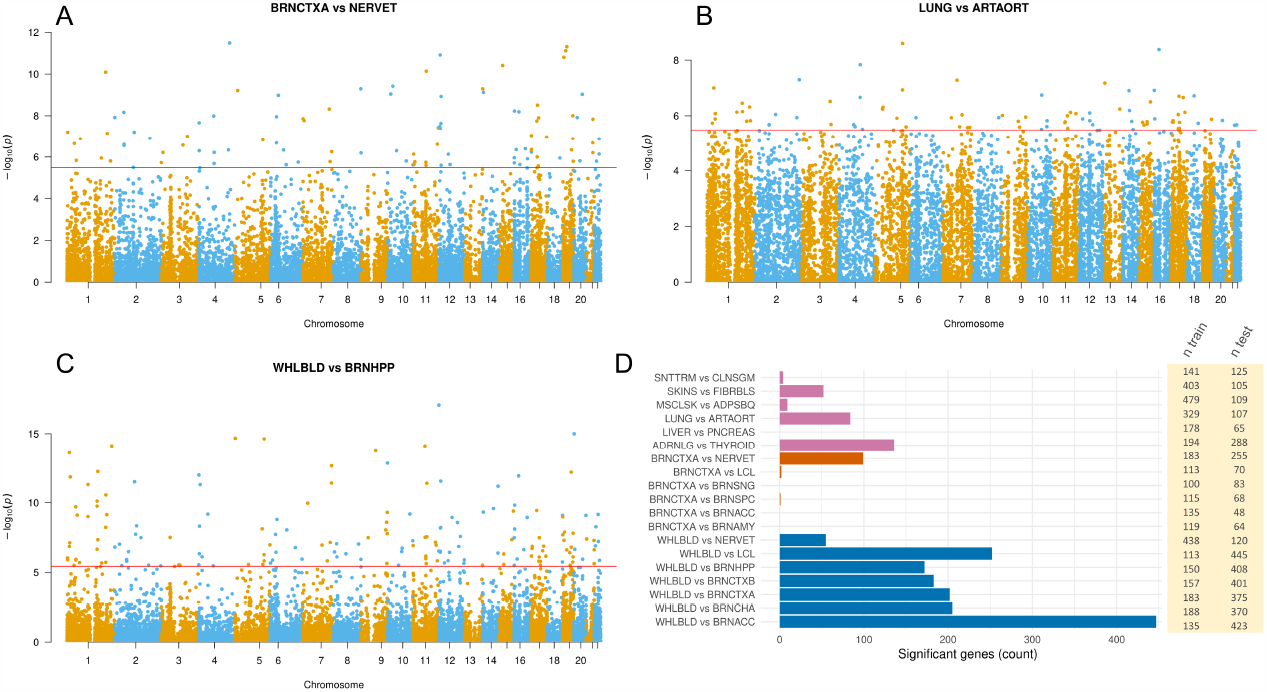
(A) (C) Manhattan plots of DRAB test results for pairings of various tissues. The horizontal red line denotes transcriptome-wide significance according to a Bonferroni multiple testing correction. (D) Numbers of differentially regulated protein-coding genes for each of 19 different tissue pairs. Statistical significance was determined according to a transcriptome-wide Bonferroni correction with a 95% confidence level.

Notably, our results confirm that biologically related tissues have more similar patterns of genetic regulation. It has been experimentally established that differences in genetic regulation, and the subsequent over- or under-expression of certain genes, are the primary drivers of tissue differentiation in animals. Consequently, tissues with similar histological structure and function are expected to exhibit similar patterns of genetic regulation. The DRAB test concurs. As shown in Figure 3, pairs of closely related brain regions have few if any genes with significantly different regulatory profiles. When comparing blood and brain tissues, on the other hand, hundreds of genes display strong evidence of tissue-specific genetic regulation. This trend also holds for the miscellaneous tissue pairs that we considered, supporting the biological reasonableness of DRAB results. Interestingly, the greatest number of differentially regulated genes, 447 in total, was identified between whole blood and Brodmann’s area 24 of the anterior cingulate cortex (BRNACC). The anterior cingulate cortex has a unique cell type profile compared to other brain regions and has been implicated in multiple psychiatric and neurological conditions (Beasley et al. (2006); Allman et al. (2011); Barthas et al. (2015); Gefen et al. (2018)), suggesting that DRAB can point to genes involved in BRNACC-specific functional pathways. It is likely, however, that our results are also impacted by the available sample sizes. Some of the tissue pairs considered had much smaller sample sizes than others, limiting the power of DRAB to detect differentially regulated genes.

## 4. Discussion

In this paper we have introduced DRAB (Differential Regulation Analysis by Bootstrapping) and described an R implementation of the method, which is available as open-source software on GitHub (https://github.com/MykMal/drab). DRAB is a statistical framework for identifying genes that have significantly different patterns of local genetic regulation between two tissues or other biological contexts. Our method utilizes penalized regression to learn context-specific models of the *cis*-genetic regulation of gene expression, and then applies a novel model comparison test to determine whether those models have equivalent predictive performance. The primary methodological innovations of our approach are twofold. First, we have proposed and confirmed that genetic regulatory profiles can be analyzed by training models to predict expression from eQTLs. Transcriptome imputation models are already widely used in statistical genetics, for instance as the first stage of TWAS, and our work illustrates another avenue for their application. Second, and more importantly, we have developed a new bootstrap model comparison test. The DRAB test incorporates both the variance associated with estimating the mean difference in model performance, as well as the variance associated with training the models themselves. Decomposing the variance in such a way makes it possible to compare the true, population-level models of genetic regulation despite the fact that they are inherently unknowable.

We have demonstrated that DRAB yields biologically reasonable results when applied to a variety of human tissue pairs from the GTEx Project. DRAB was able to effectively control type I error when run on a negative control created from random samples of the same tissue and was robust to a variety of simulation parameters. It also had sufficient power to detect genes that exhibit tissue-specific differences in regulatory profiles when such genes were expected to be found. Moreover, DRAB detected fewer differentially regulated genes in tissue pairs that share similar structure and function (e.g. different brain regions) than in pairs of biologically unrelated tissues (e.g. blood and brain tissues). This result is consistent with the contemporary understanding of gene expression and its relation to tissue function.

Although our application was only concerned with identifying genes that have tissuespecific patterns of genetic regulation of mRNA expression, the DRAB framework has broad relevance to many other problems in molecular genetics and applied statistics. In fact, DRAB can also test for differential genetic regulation in other gene expression products (i.e. other omic data types). The software package that we developed is directly applicable to any genetically influenced quantitative molecular phenotypes, including micro RNA expression, intron excision ratios, and measurements of transcription factor activity. More generally, our bootstrap model comparison test facilitates the rigorous assessment of predictive performance in any regression task where multiple competing parametric models are being considered. Model comparison and benchmarking are active areas of machine learning research, so we plan to further develop the ideas introduced herein.

We conclude with several limitations of our approach. First, we have not conducted a theoretical study of the proposed model comparison test. Some conditions may be necessary for asymptotic validity, such as consistency and asymptotic normality of parameter estimation with the candidate models (Politis and Romano (1994)). Since we are considering linear parametric models and the elastic net is known to have model selection consistency (Yuan and Lin (2007); Jia and Yu (2010)), we expect those conditions to hold. However, it is possible that some adjustments might be needed, such as refitting selected models by least squares without penalization (Zhao, Witten and Shojaie (2021)) or de-biasing via desparsified lasso and ridge projection procedures (Dezeure et al. (2015)). More generally, the proposed method may or may not perform well with non-parametric models due to technical challenges (Dai, Shen and Pan (2022)). Second, DRAB does not estimate a meaningful effect size for differential regulation. It can identify the genes that are differentially regulated between any two biological sample types, but it cannot determine whether some of those genes exhibit more extensive differences in genetic regulation than the others. For the same reason, DRAB is also unable to reconstruct the regulatory networks that underlie its results. We suggest possibly using DRAB in conjunction with differential network inference methods, which aim to build context-specific regulatory networks but cannot perform tests of statistical significance (Bhuva et al. (2019); Baur et al. (2020); Yazdani et al. (2020)). Third, the utility of DRAB ultimately depends on its ability to learn accurate models of transcriptional regulation. A similar limitation exists in TWAS, which is known to lose power when the *R*^2^ of transcriptome imputation models is low (He et al. (2022)). In future work it may be beneficial to evaluate how the choice of prediction algorithm affects the power of DRAB to detect differentially regulated genes. Lastly, a major practical downside of our method is that it requires individual-level genotypic and phenotypic data. Extensions of DRAB to genome-wide association study (GWAS) summary data would greatly expand the scope of its applications in genetics.

## Acknowledgments

We thank the associate editor and two anonymous reviewers who provided very thoughtful and constructive feedback on the initial version of this manuscript. Access to the GTEx data was approved for dbGaP Project #26511, and the data were obtained from dbGaP accession number phs000424.v8.p2 on 10/13/2021. We are also grateful to the Minnesota Supercomputing Institute (MSI) at the University of Minnesota for providing resources that contributed to the results reported within this paper.

## Funding

MMM was supported by NIH grant T32GM132063; XTS and WP were supported by NIH grants U01AG073079, R01AG065636, R01AG069895, R01AG074858, and R01GM126002. The content is solely the responsibility of the authors and does not necessarily represent the official views of the National Institutes of Health.

## SUPPLEMENTARY MATERIAL

### Additional figures for

This PDF file contains additional figures with results from applications of DRAB to simulated and real data. In particular, we provide figures showcasing the results of sensitivity analyses and QQ plots of DRAB *P* -values for every tissue pair analyzed in the main text.

### Source data for

This ZIP archive contains tab-separated value (TSV) files with all of the numerical results from our application of DRAB to GTEx data. Included are the DRAB test *P* -values, conditional *t*-test *P* -values, and training/test set sample sizes for every tissue pair and gene.

## Notes

https://github.com/MykMal/drab

### Competing Interest Statement

The authors have declared no competing interest.

### Summary of Updates

New section describing simulation studies; new supplementary file with additional figures; improvements to text throughout.

