## Additional figures for "A bootstrap model comparison test for identifying genes with context-specific patterns of genetic regulation"

#### 1 Simulation results for null comparisons

Below are quantile-quantile (QQ) plots of $P$-values from the DRAB differential regulation test applied to simulated gene expression data and real genotype data from the GTEx v8 data set. Comparisons are between independent simulations of the same tissue, e.g. testing for differential regulation between one simulated brain cortex data set and another simulated brain cortex data set. Training set sample sizes were fixed at 238 and test set sample sizes were fixed at 239, while the choice of test set tissue and the amount of sample overlap between training sets were varied. The same 50 autosomal protein-coding genes were included in each simulation study. The theoretical null distribution (assuming no data set-specific genetic regulation) is plotted as a solid gray line, flanked by 95% confidence bands.

##### 1.1 Trained and tested on whole blood (WHLBLD)

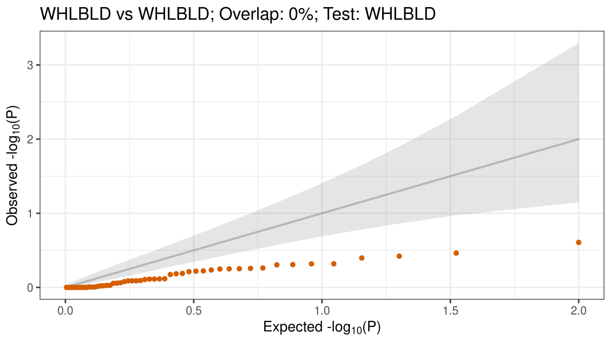


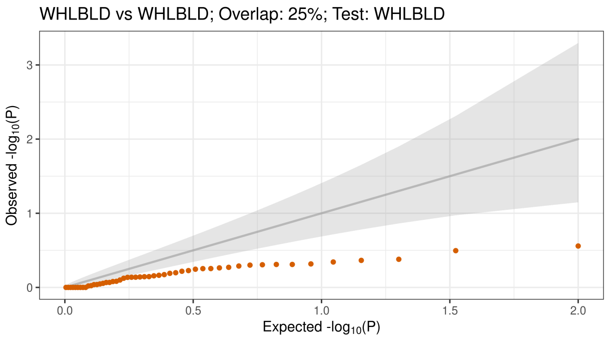


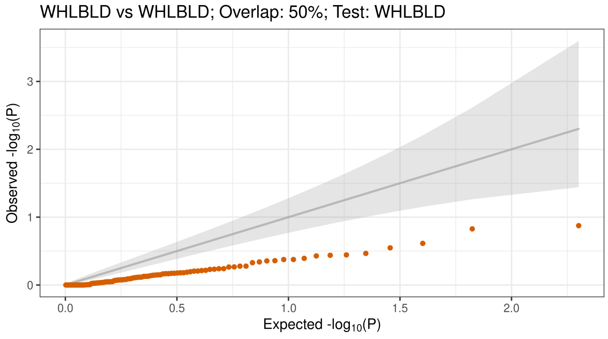


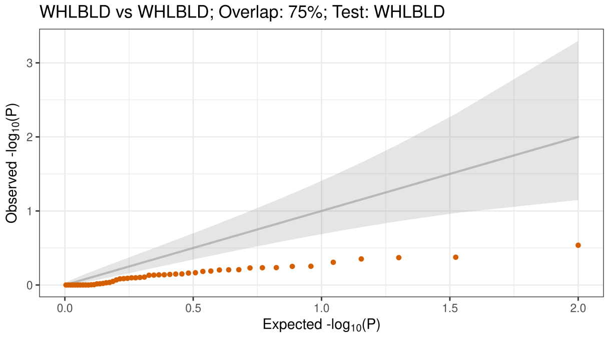

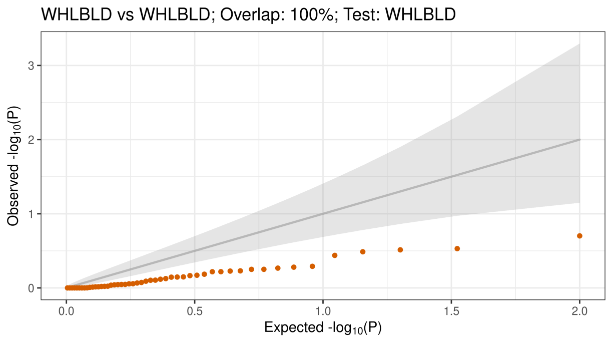


#### 1.2 Trained and tested on brain cortex (BRNCTXA)

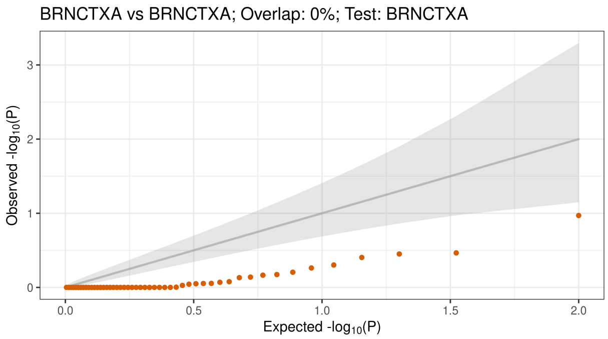


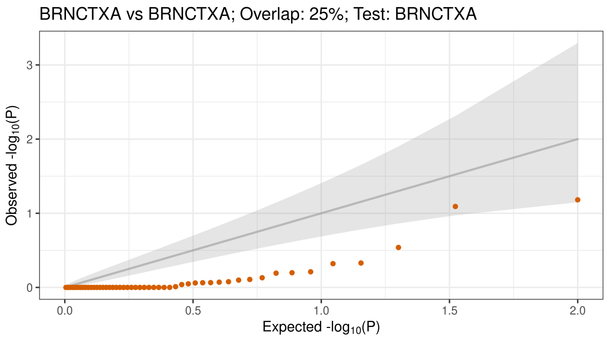


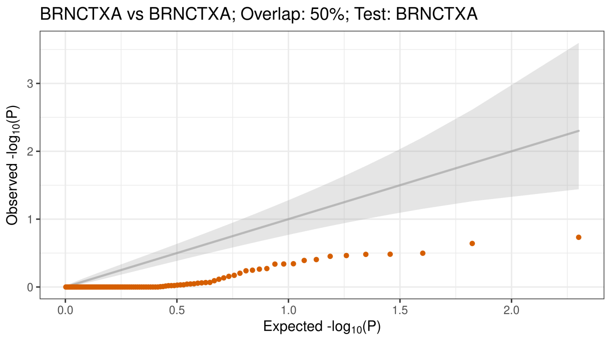


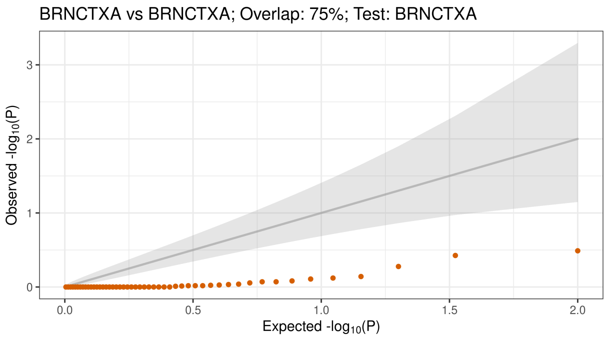


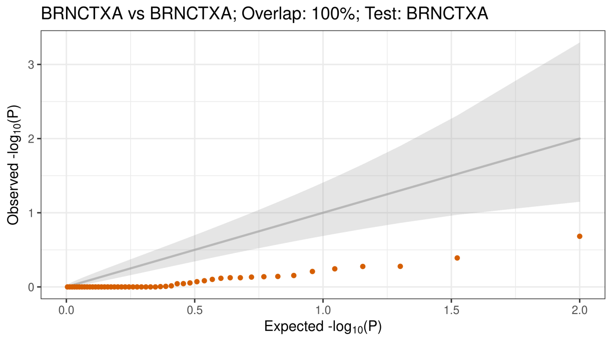

#### 1.3 Trained on brain cortex (BRNCTXA), tested on whole blood (WHLBLD)

BRNCTXA vs BRNCTXA; Overlap: 0%; Test: WHLBLD

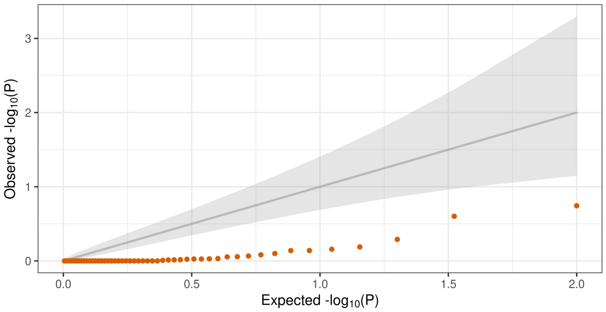


BRNCTXA vs BRNCTXA; Overlap: 25%; Test: WHLBLD

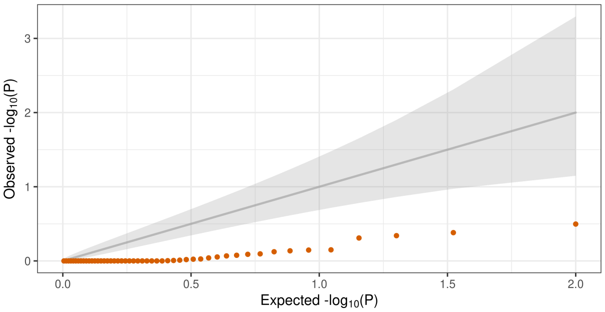


BRNCTXA vs BRNCTXA; Overlap: 50%; Test: WHLBLD

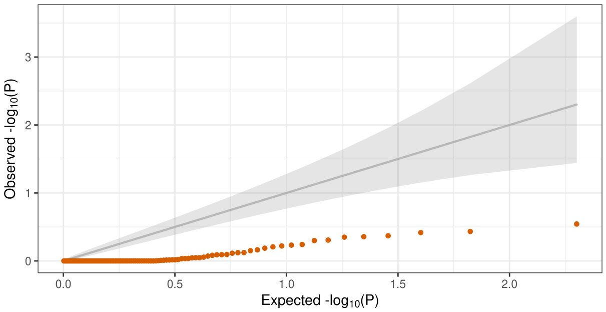


BRNCTXA vs BRNCTXA; Overlap: 75%; Test: WHLBLD

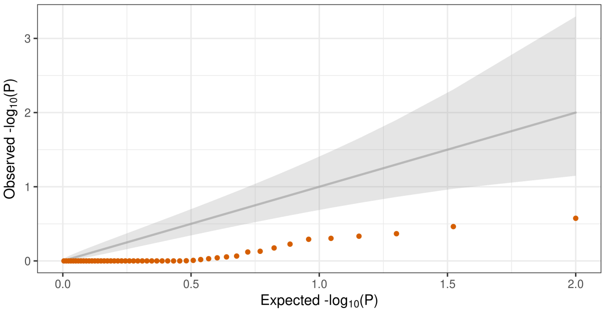


BRNCTXA vs BRNCTXA; Overlap: 100%; Test: WHLBLD

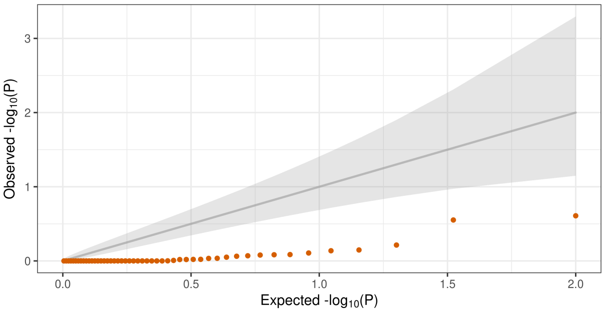


#### 1.4 Trained on whole blood (WHLBLD), tested on brain cortex (BRNCTXA)

WHLBLD vs WHLBLD; Overlap: 0%; Test: BRNCTXA

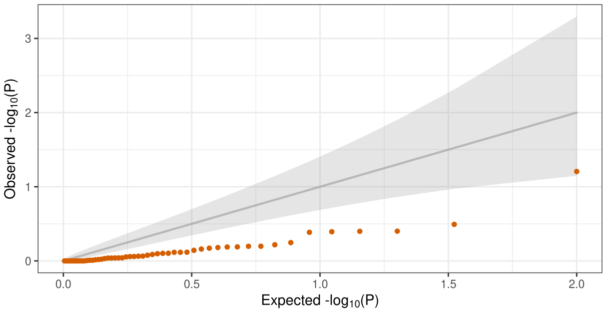


WHLBLD vs WHLBLD; Overlap: 25%; Test: BRNCTXA

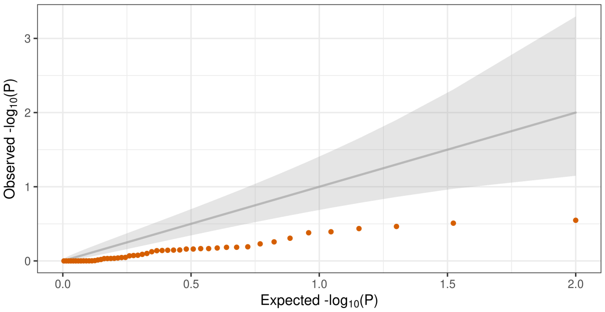

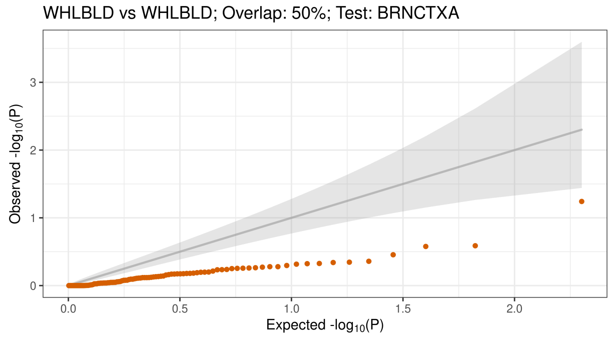


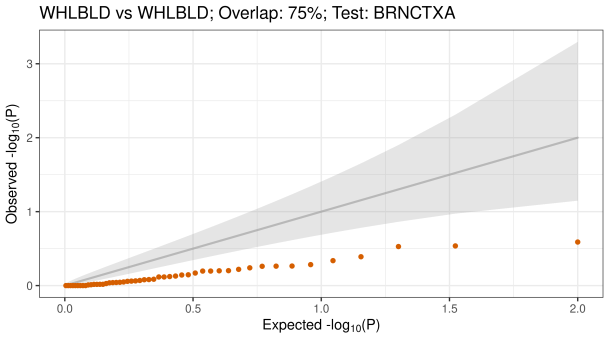


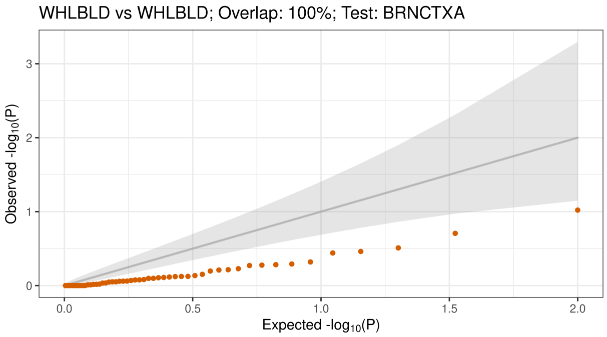


### 2 Simulation results for power analyses

Below are scatter plots of $P$-values from the DRAB differential regulation test applied to simulated gene expression data and real genotype data from the GTEx v8 data set. Each comparison is between simulated whole blood expression data and simulated brain cortex expression data. All simulation parameters except those specified in the plot labels were fixed at the following default values: each training set had a sample size of 238, there was no sample overlap between the two training sets, and the test set had a sample size of 239. The same 50 autosomal protein-coding genes from the previous section were included in each simulation study.

#### 2.1 Effect of sample overlap when testing on brain cortex (BRNCTXA)

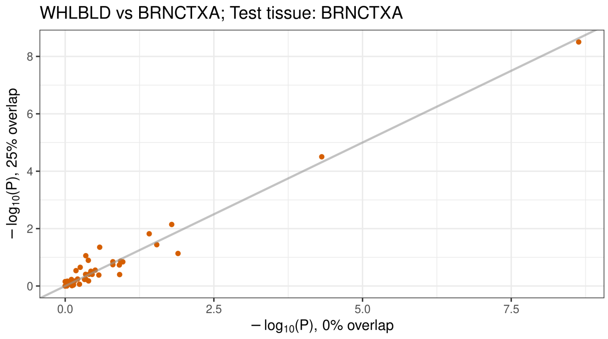


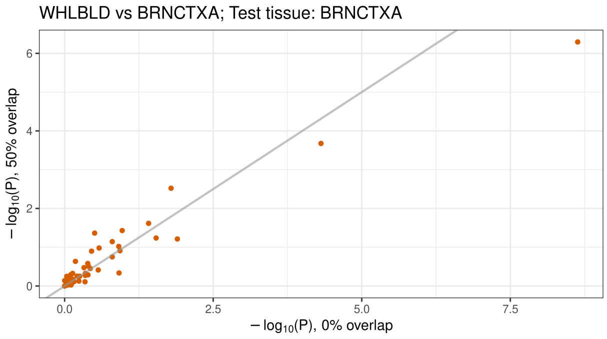

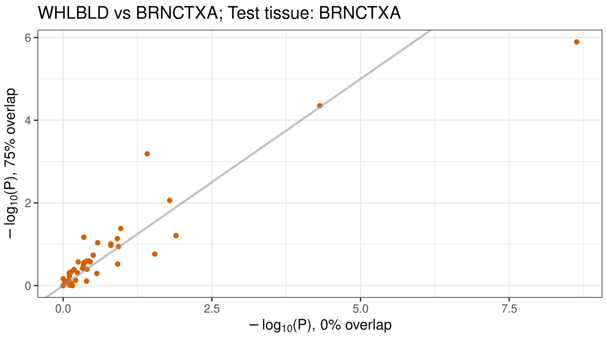


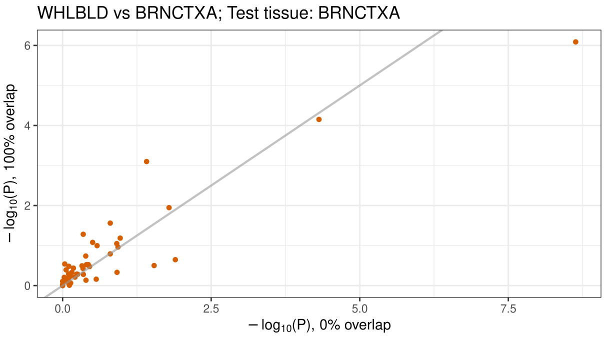


#### 2.2 Effect of sample overlap when testing on whole blood (WHLBLD)

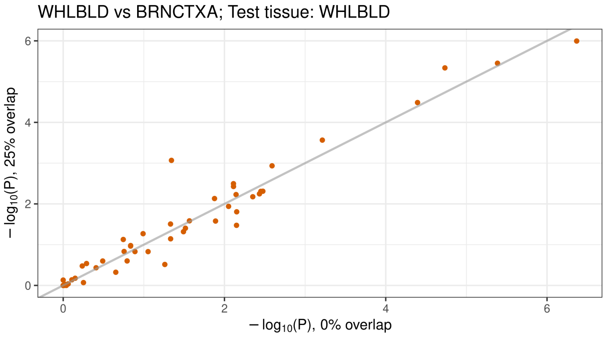


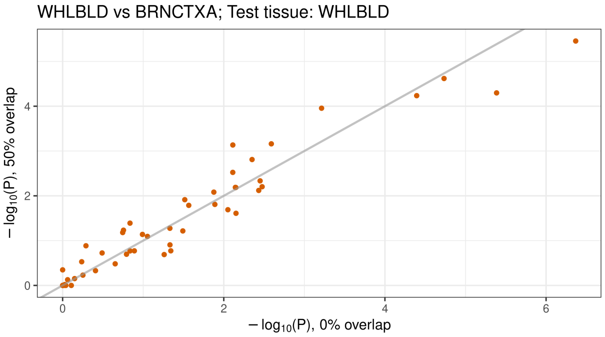


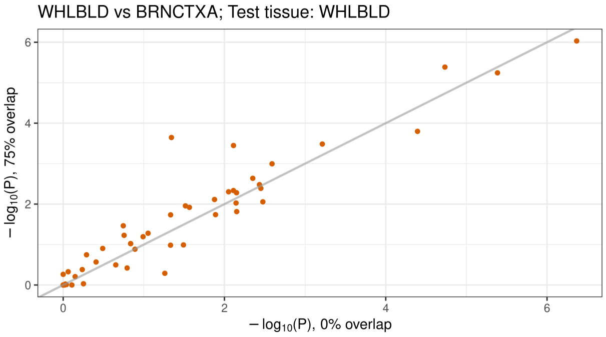


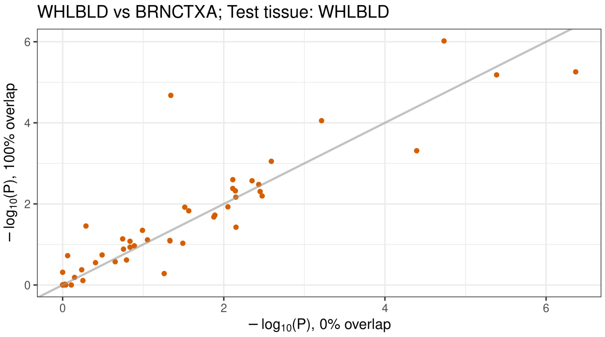


#### 2.3 Effect of test tissue choice

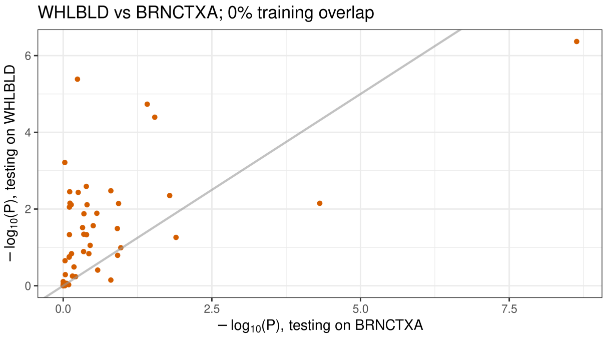


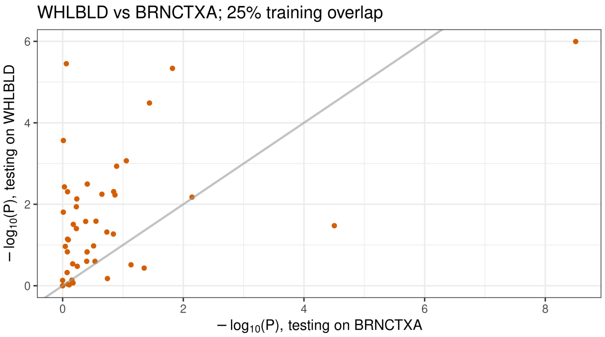

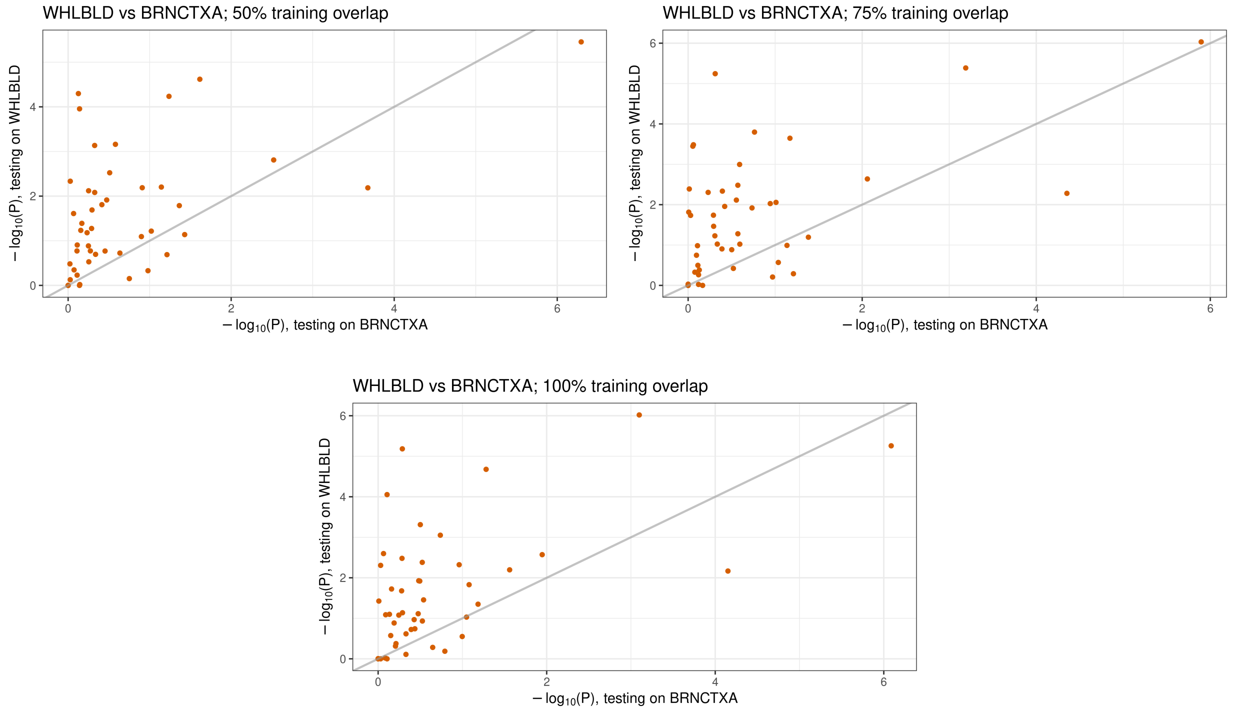


#### 2.4 Effect of test set sample size when testing on whole blood (WHLBLD)

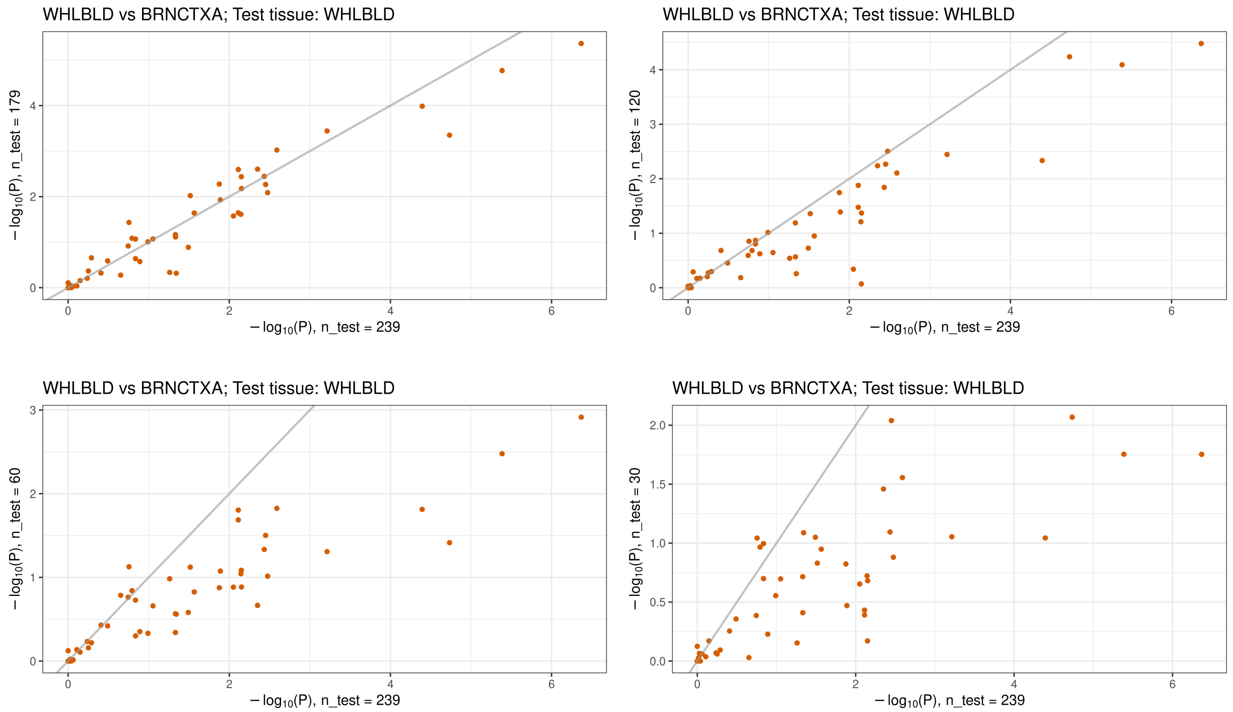

#### 2.5 Effect of test set sample size when testing on brain cortex (BRNCTXA)

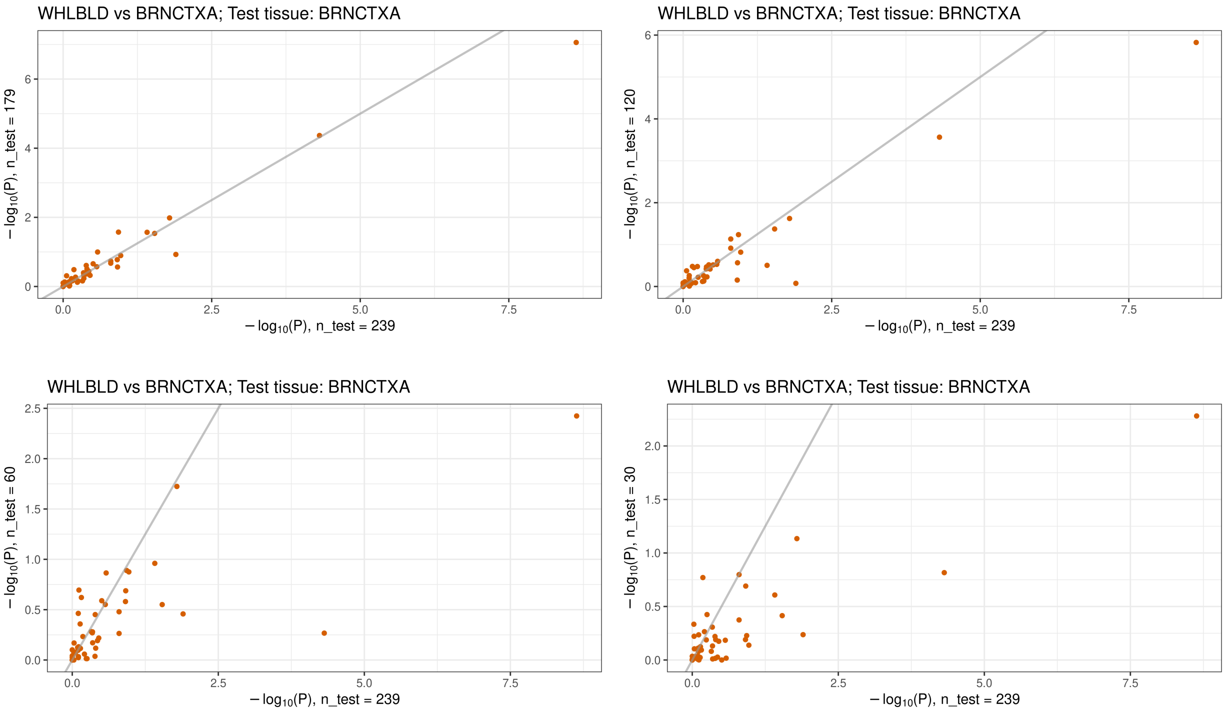


#### 2.6 Effect of training set sample size

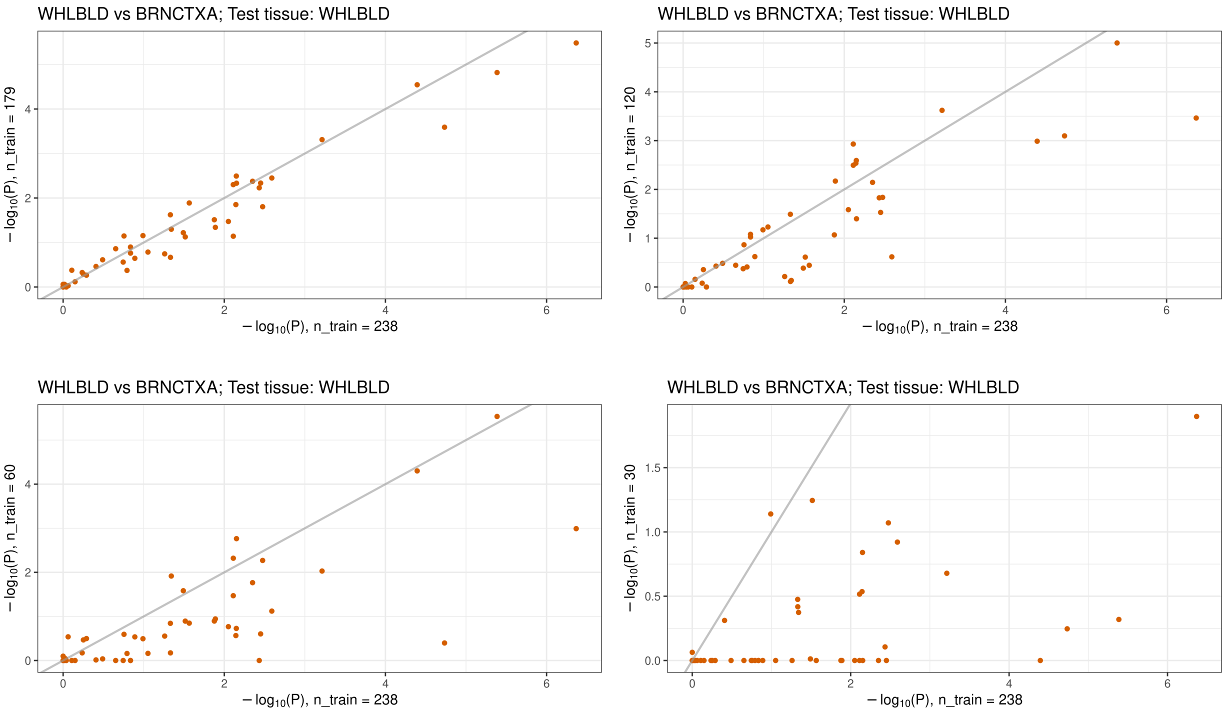

### 3 QQ plots for all real data tissue pairs

Below are quantile-quantile (QQ) plots of $P$-values from the DRAB differential regulation test for every tissue pair considered in our real data analysis. All autosomal protein-coding genes were included in the tests. The theoretical null distribution (assuming no tissue-specific genetic regulation) is plotted as a solid gray line, flanked by 95% confidence bands. Definitions for all tissue abbreviations are provided in Table 1 in the main text.

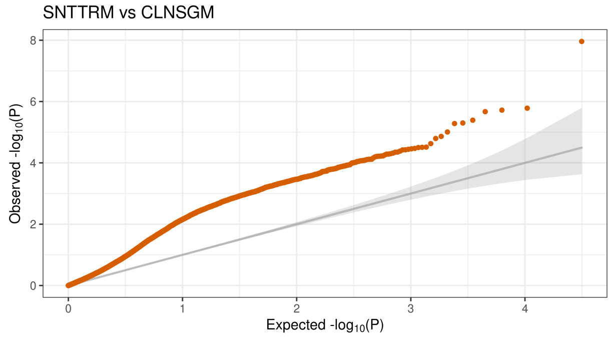


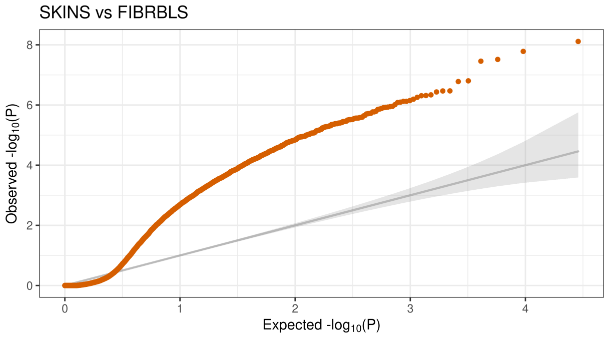


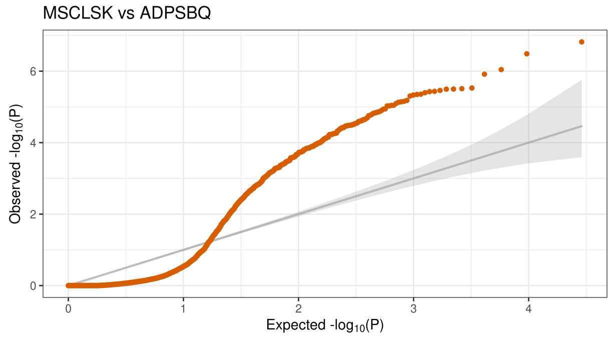


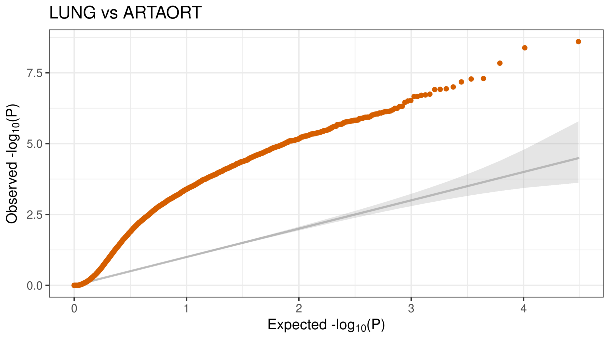


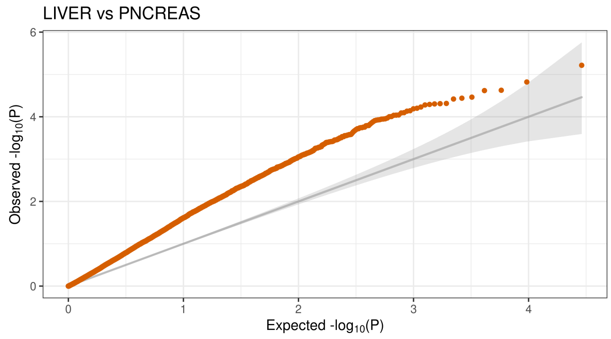


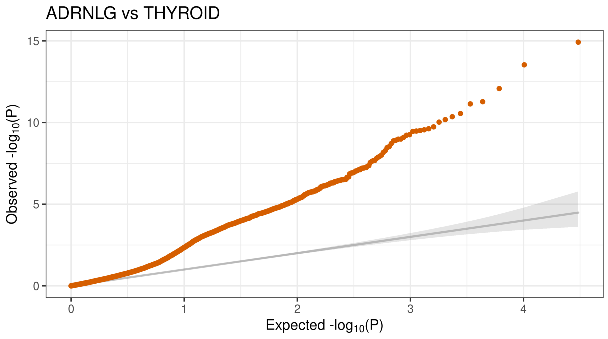

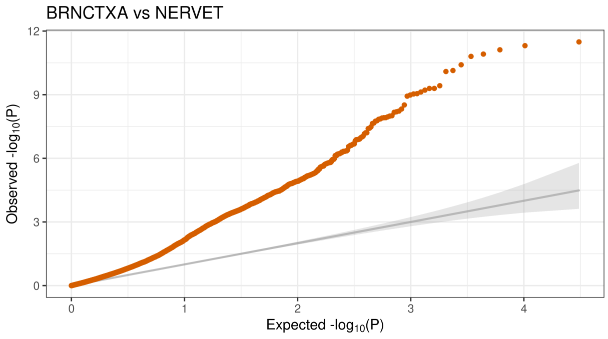
BRNCTXA vs NERVET
Observed -log10(P)
Expected -log10(P)

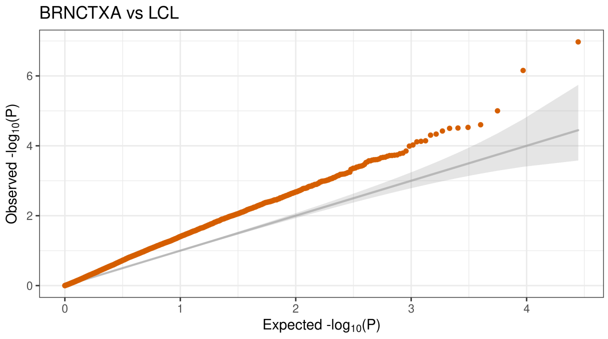
BRNCTXA vs LCL
Observed -log10(P)
Expected -log10(P)

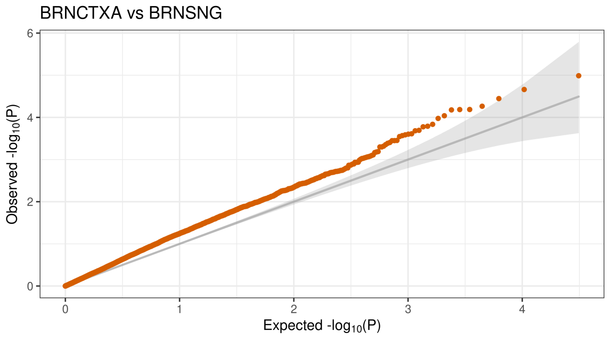
BRNCTXA vs BRNSNG
Observed -log10(P)
Expected -log10(P)

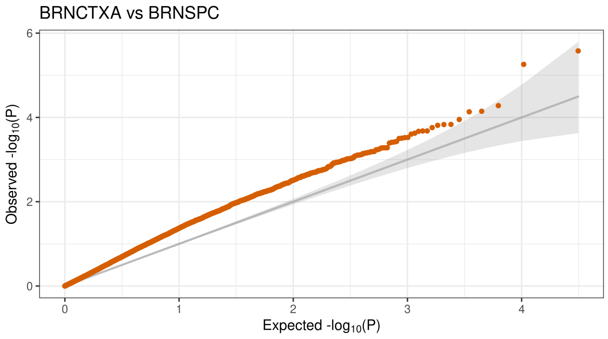
BRNCTXA vs BRNSPC
Observed -log10(P)
Expected -log10(P)

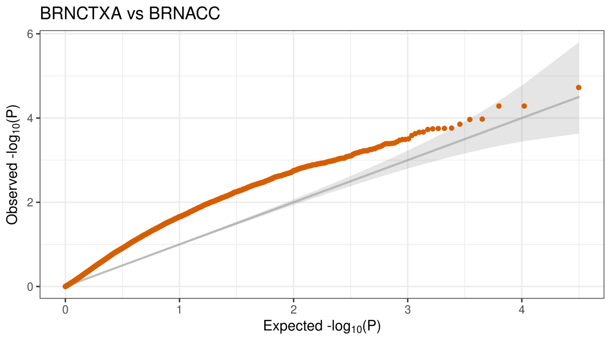
BRNCTXA vs BRNACC
Observed -log10(P)
Expected -log10(P)

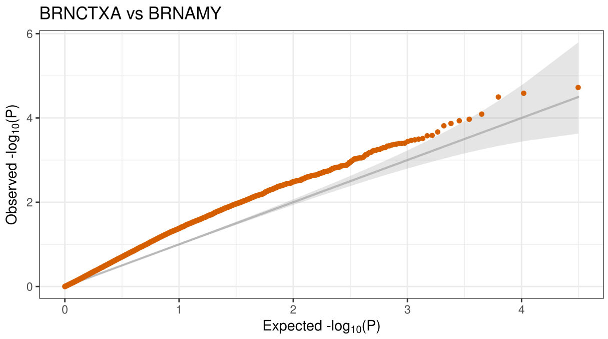
BRNCTXA vs BRNAMY
Observed -log10(P)
Expected -log10(P)

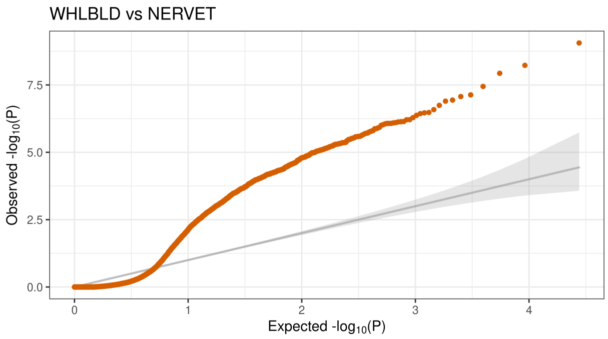
WHLBLD vs NERVET
Observed -log10(P)
Expected -log10(P)

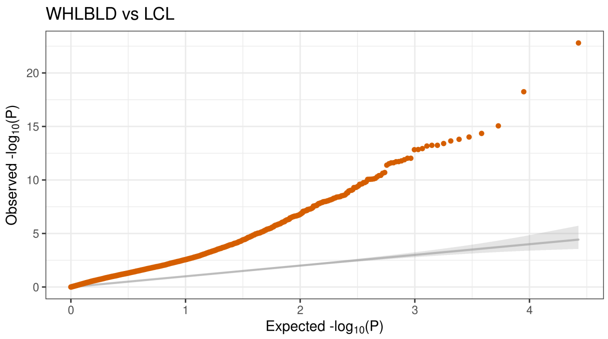
WHLBLD vs LCL
Observed -log10(P)
Expected -log10(P)

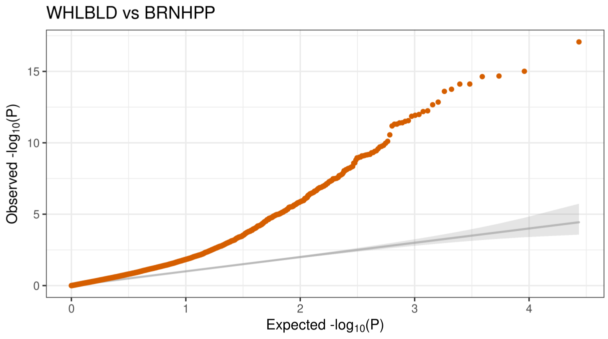
WHLBLD vs BRNHPP
Observed -log10(P)
Expected -log10(P)
0
5
10
15
0
1
2
3
4

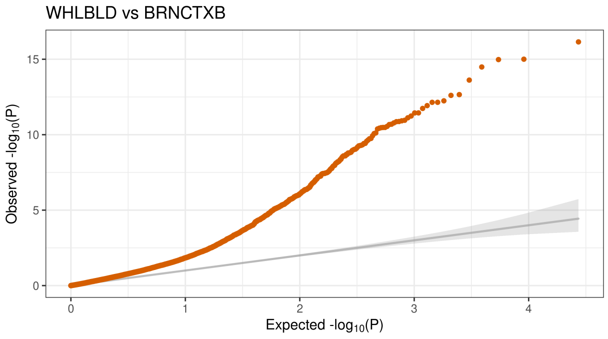
WHLBLD vs BRNCTXB
Observed -log10(P)
Expected -log10(P)
0
5
10
15
0
1
2
3
4

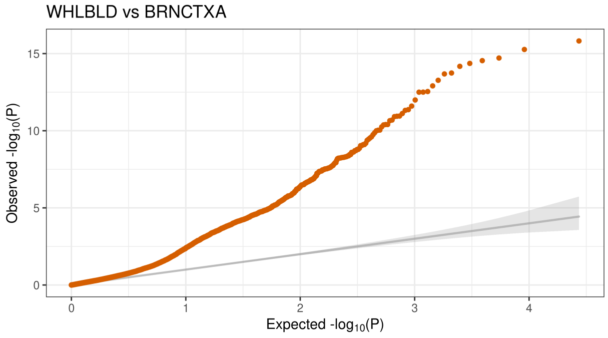
WHLBLD vs BRNCTXA
Observed -log10(P)
Expected -log10(P)
0
5
10
15
0
1
2
3
4

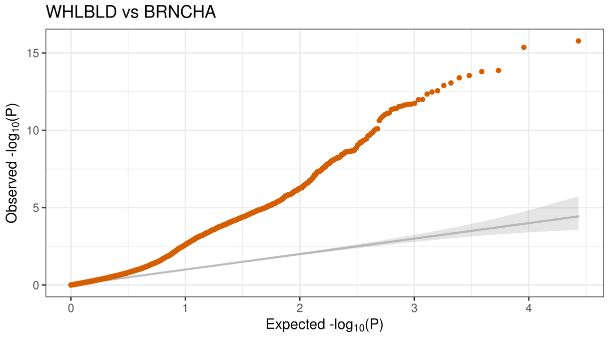
WHLBLD vs BRNCHA
Observed -log10(P)
Expected -log10(P)
0
5
10
15
0
1
2
3
4

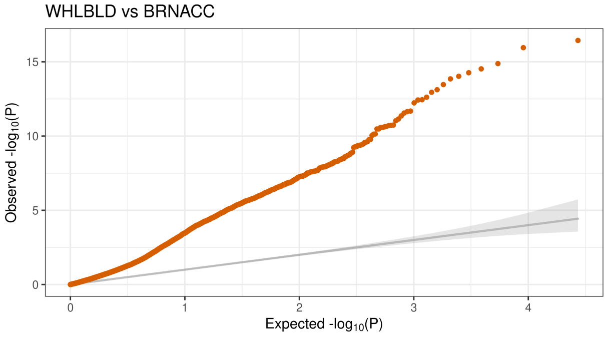
WHLBLD vs BRNACC
Observed -log10(P)
Expected -log10(P)
0
5
10
15
0
1
2
3
4
